# State and trait serotonin variations interact to shape the intrinsic connectivity and gradient architecture of the brain – a combined TPH2 genetics and tryptophan depletion study

**DOI:** 10.1101/2024.09.29.615637

**Authors:** Lan Wang, Congcong Liu, Ting Xu, Xianyang Gan, Keith Kendrick, Weihua Zhao, Christian Montag, Benjamin Becker

**Affiliations:** The Center of Psychosomatic Medicine, Sichuan Provincial Center for Mental Health, Sichuan Provincial People’s Hospital, University of Electronic Science and Technology of China, Chengdu, China; School of Life Science and Technology, University of Electronic Science and Technology, Chengdu, China; School of Psychology, Xinxiang Medical University, Xinxiang, China; Faculty of Psychology, Southwest University, Chongqing, China; Key Laboratory of Cognition and Personality, Ministry of Education, Chongqing, China; Department of Psychology, Ulm University, Ulm, Germany; Department of Psychology, The University of Hong Kong, Hong Kong, China; State Key Laboratory of Brain and Cognitive Sciences, The University of Hong Kong, Hong Kong, China

**Author notes:** Correspondence Benjamin Becker, The University of Hong Kong, Department of Psychology. Shared first-authorship (Lan Wang, Congcong Liu, Ting Xu).

**Keywords:** Serotonin, tryptophan depletion, Tryptophan hydroxylase, vmPFC, amygdala, connectome gradient

## Abstract

**Background:** Serotonin (5-HT) critically regulates cognitive and emotional functions, and both stable and transient variations in 5-HT signaling have been associated with emotional dysregulations. However, findings regarding the neurofunctional effects of transient 5-HT variations have been highly inconsistent. Therefore, we examined whether individual variations in a central 5-HT-regulating genetic polymorphism (tryptophan hydroxylase 2, TPH2) represent a vulnerability or resilience factor for the effects of acute tryptophan depletion (ATD) on functional brain architecture.

**Method:** The current study utilized a pharmacogenetic within-subject randomized placebo-controlled resting-state fMRI design with n=53 healthy male participants in combination with spontaneous intrinsic neural activity, functional connectivity, and connectome gradient analyses to compare the neurofunctional effects of ATD-induced transient reduction in central 5-HT signaling between TPH2 genotypes (a priori genotyping for rs4570625, GG n = 25 vs. TT n = 23).

**Results:** ATD induced significant increases in spontaneous neural activity in hippocampal CA1 irrespective of genotype and enhanced communication of this region with the bilateral amygdala and the vmPFC specifically in GG carriers. ATD sharpened the intrinsic connectome gradient architecture in several large-scale networks, including the salience, frontoparietal, and default mode network.

**Conclusions:** Our results identify a potential genetic marker for an increased vulnerability to the neural effects of transient variations in 5-HT signaling on the functional architecture of an anxiety- and stress-related brain circuit. Connectome gradient results underscore the regulatory role of 5-HT on the intricate organization of large-scale networks involved in emotional reactivity and regulation.

## Introduction

Serotonin (5-HT) is a neurotransmitter synthesized from the amino acid tryptophan in the dorsal and median raphe nucleus (1) and plays an important role in a range of physiological as well as cognitive and affective functions, including executive functions, emotion and emotion regulation (2, 3, 4, 5). The cognitive and affective effects are mediated via dense ascending projections from the dorsal raphe to the limbic system, including the amygdala and hippocampus, and nearly the entire frontal cortex.

Dysfunction in serotoninergic neurotransmission in these pathways has been considered as candidate mechanisms underlying emotional dysregulations in response to stress, impulsive and aggressive behavior and affective dysregulations in anxiety disorders (6, 7). Preclinical studies have further underscored the role of limbic structures, including the amygdala and hippocampus and the prefrontal cortex (8) as pivotal sites of dense serotoninergic innervation (9) as well as in mediating the function of 5-HT in emotional reactivity (10), stress and emotion regulation (11, 12, 13).

5-HT signaling is influenced by stable, trait-like, individual differences, including genetics (e.g., 14) and by transient variations in the production of 5-HT in the brain (e.g., influenced by stress or nutrition, (15, 16). The putative role of 5-HT in emotional processes has been extensively investigated using acute tryptophan depletion (ATD) protocols (17), a procedure leading to a transient reduction in central serotonin levels. ATD entails the depletion of the serotonin amino acid precursor – tryptophan - thereby reducing central serotonin synthesis, resulting in a transient decrease in central tryptophan and serotonergic signaling (18). The procedure has been successfully employed to explore the role of 5-HT in cognitive and emotional functions (19, 20) and its interaction with other signaling systems (e.g., oxytocin, 3, 13). Rodent models have demonstrated that ATD modulates 5-HT concentrations in the cortex and limbic system (21) while ATD-induced reduced availability of 5-HT signaling in humans elevated amygdala responsiveness to negative emotional stimuli including angry (22) and fearful (23) faces and enhanced amygdala-medial prefrontal cortex resting-state coupling (24) highlighting the modulatory influence of 5-HT and limbic-prefrontal circuits involved in threat reactivity and emotion regulation (2, 25, 26, 27). Participants undergoing ATD exhibited altered responses to negative emotional stimuli and aggressive provocation (28) underpinned by modulation of the ventromedial prefrontal cortex (vmPFC) (29, 30) and amygdala connectivity (22). Cools et al. additionally reported that during processing of emotional signals ATD modulated the amygdala-hippocampal interactions (31), a pathway previously associated with the level of stress exposure (13). ATD-induced manipulation of serotonergic levels also to modulations in the amygdala-hippocampus and amygdala-prefrontal circuits probably mediating sensitivity to negative emotional states and stress exposure (32, 33).

However, recent meta-analytical studies did not find strong evidence for robust effects of ATD on emotional and neural processing in healthy individuals (34, 35) suggesting that its impact on emotion and impulsive behavior is notably limited compared to the robust effects on the neurochemical level (36). While these findings challenge a series of observations on ATD-induced emotional and neural changes, accumulating findings from recent studies suggest considerable individual variations in the effects of ATD and that these vary as a function of trait-like and genetic factors which may mirror individual variations in the basal function and architecture of the monoamine, including the 5-HT system (e.g.,14, 37, 38) and polymorphisms in the tryptophan hydroxylase 2 (TPH2) gene (4) The TPH2 encodes the rate-limiting enzyme responsible for central serotonin synthesis from tryptophan and individual differences in the TPH2 have been associated with serotonin synthesis rates in limbic and prefrontal brain regions (39, 40, 12), alongside variations in emotional responsiveness and regulation (41, 42). Specifically, the TPH2 rs4570625 single nucleotide polymorphism (SNP), situated within the gene’s promoter region (43), with variations on this SNP being associated with individual differences in the intrinsic functional organization of the brain, including variations in amygdala functional connectivity (12) and macroscale whole-brain structural connectivity (44). Consequently, variations in TPH2 rs4570625 SNP may underlie individual differences in the functional communication of critical emotion processing hubs in the brain and may furthermore mediate individual differences in the sensitivity of the effects of transient decrease in 5-HT signaling via ATD on these networks.

To this end the current study utilized a pre-registered within-subject randomized placebo-controlled double-blind resting state fMRI (rsfMRI) design in a sample of n=53 healthy male participants (design details see also 4) to explore the underlying regulatory mechanisms of TPH2 rs4570625 SNP on the neurofunctional sensitivity to an ATD-induced transient reduction of central 5-HT signaling, specifically on the vmPFC-hippocampus-amygdala circuitry and whole brain gradient architecture of the brain. To determine the interactions between ATD and genotype extreme groups of the TPH2 rs4570625 SNP were sampled (TT vs GG). We initially conducted a voxel-wise fractional amplitude of low-frequency fluctuations (fALFF) analysis -a measure closely linked to spontaneous intrinsic neural activity and with a high sensitivity to metabolic activity and pharmacological receptor engagement (e.g., 45, 46) – to determine regions sensitive to 5-HT variations with a high spatial resolution. Next, network level changes were examined using seed-based voxel-wise functional connectivity (FC) analysis to further identify interactions between genetic variation and acute depletion of 5-HT. Finally, we employed a connectome gradient analysis to determine variations in large-scale subnetwork architecture. We hypothesized that ATD-induced reduction in 5-HT signaling would (1) evoke regional-specific changes in spontaneous neural activity and connectivity in the vmPFC-hippocampus-amygdala systems; and (2) modulate the large-scale gradient architecture of the brain; and that (3) these neural effects would vary as a function of TPH2 gene polymorphism.

## Methods

### Participants

53 right-handed, healthy male participants were recruited in total. To reduce variance related to sex differences, e.g., related to sex differences in central 5-HT synthesis rates and the cognitive effects of ATD (47, 48, 49), the present study recruited male individuals only (3, 13). Details in the procedures are described in our previous task-based fMRI study on threat reactivity using the same protocols and sample (4). Please note that the task-paradigm followed the resting state acquisition reported here, and two participants only underwent the resting state scan due to the decision not to participate in a threat paradigm (sample size in the present study n = 53, previous n = 51). All participants were free from a current or a history of medical, neurological or psychiatric disorder, regular use of psychotropic substances or MRI contraindications. Additionally, 5 subjects were excluded due to excessive head movement (>2mm translation or 2° rotation, n = 3) and technical failure of the MRI system (n = 2), leading to a final sample of n = 48 participants for the current analyses (age range 18-27 years, mean age = 21.94 ± 2.59 years, n = 23 TT carriers). In line with the design of our previous study Liu et al., 2023 (Ref 4) genotypic extreme groups were chosen, such that a comparable large number of TT vs. GG participants (23 vs. 25) were recruited from a genetic data bank. This approach helps to increase the statistical power while testing for genetic effects of rs4570625 or interaction with treatment effects, respectively. Details on the recruitment procedures and genotyping procedures are provided in our previous study by Liu et al., 2023 (Ref 4). The present manuscript reports the findings from the resting state fMRI assessment. The studies were approved by the local ethics committee and in accordance with the latest revision of the Declaration of Helsinki. All subjects provided written informed consent after being fully informed about the experimental procedures. Study protocols were pre-registered on clinicaltrials.gov(https://clinicaltrials.gov/ct2/show/NCT03549182, ID NCT03549182).

### Assessment of potential confounders

Given that emotional state and childhood experience can influence neural processing or central 5-HT signaling and can interact with TPH2 genetics (50, 51, 52, 53, 12), trait and state anxiety (State-Trait Anxiety Inventory, STAI, 54), current stress (Perceived Stress Scale, PSS, 55) and exposure to early life stress (Childhood Trauma Questionnaire, CTQ, 56) were assessed before treatment administration to control for differences between the genotype groups. The validated Chinese versions of these assessment were utilized in the current study (57, 58, 59).

### Procedure

Experimental and treatment procedures were identical to our previous study that reported the results of the task-based (threat exposure) fMRI paradigm that followed the resting state fMRI assessment (4). The studies aimed at determining whether TPH2 genotype can render individuals vulnerable to the neurofunctional effects of acute tryptophan depletion during threat exposure (published as 4) or on the intrinsic brain functional architecture. To this end, extreme groups of the TPH2 s4570625 were recruited (GG vs TT carriers) and underwent a within-subject randomized double-blind placebo-controlled pharmaco-rsfMRI design. Eligible participants were administered an oral amino acid mixture (ATD, total weight, 100g), which causes a transient decrease in central 5-HT levels, or a control placebo drink (total weight, 102.3g) in a randomized order during two experimental sessions. Participants were instructed to abstain from alcohol and caffeine for 24h and from food and drinks (except water) for 12h prior to the experiment. To adhere to the pharmacodynamic profile of treatment, participants arrived between 7:30 to 10:00 AM and underwent fMRI acquisition between 13:00 to 15:30 PM. Upon arrival, participants received a standardized protein-poor diet for breakfast. Participants next underwent a previously validated tryptophan depletion protocol with ATD which has been demonstrated to lead to a robust transient reduction in central 5-HT signaling (60, 61, 62, 63) or PLC. Previous studies have shown that administration of an ATD causes a sustained decrease in serotonin levels, with the effect reaching a maximum after approximately 5 hours and lasting for 2 hours (64, 65, 66). Therefore, participants underwent rsfMRI acquisition 5 hours after treatment administration (duration 7.5 minutes) followed by the task-based fMRI paradigms. All subjects received both ATD and PLC treatments with an interim wash-out period of approx. 5 weeks.

### MRI data acquisition

MRI data were acquired on a 3.0 Tesla GE MR750 system (General Electric Medical System, Milwaukee, WI, USA). To exclude subjects with apparent brain pathologies and improve normalization of the functional times series, T1-weighted high-resolution anatomical images were acquired with a spoiled gradient echo pulse sequence: repetition time (TR) = 5.9ms; flip angle = 9°; field of view (FOV) = 256 × 256mm; acquisition matrix = 256 × 256; slice thickness = 1mm; number of slices = 156. Restingstate data fMRI data were acquired using an T2*-weighted echo planar imaging sequence utilizing following parameters: TR = 2000ms; TE = 30ms; FOV = 220 × 220mm; flip angle = 90°; image resolution = 64 × 64; thickness = 3.2mm.

### MRI data preprocessing

All MRI data were preprocessed using the GRETNA package (https://www.nitrc.org/projects/gretna,67) implemented in MATLAB R2014a. A standardized preprocessing pipeline referred to GRETNA manual (68) and was employed and included discarding the first volumes acquired during the initial 10 seconds for each participant, slice timing correction, realignment, spatial normalization to the Montreal Neurological Institute (MNI) space, spatial smoothing with a Gaussian Kernel of 6mm full width at half maximum (FWHM), temporally linear detrending to diminish the effects of drift or trends in the signal, nuisance regression, temporal filtering (0.01-0.1 Hz) and scrubbing to reduce the effects of head motion.

### Region of interest (ROI) definition

Based on our regional specific hypotheses based on prior studies our analyses focused on the hippocampus, amygdala and vmPFC. In accordance with our previous study the hippocampus and amygdala ROIs were derived from Automatic Anatomical Labelling (69). The vmPFC was defined based on an automated analysis of 199 studies using NeuroSynth (www.neurosynth.org) with the search term “vmPFC” and thresholded at p<0.05 with false discovery rate correction (FDR, i.e. p_FDR_<0.5) (4).

### Voxel-wise fractional amplitude of low-frequency fluctuations (fALFF) analysis

The fractional amplitude of low-frequency fluctuations (fALFF) quantifies the relative contribution of frequency fluctuations within a specific frequency band (4) to the whole detectable frequency range (70) which is closely related to regional spontaneous neuronal activity(71). Moreover, fALFF has shown a robust replicability of pharmacological modulation of spontaneous brain activity (46). To initially examine effects of genotype and acute 5-HT reduction on spontaneous neural activity we initially calculated fALFF maps for each subject based on unfiltered preprocessed data. Next contrast maps between ATD and PLC treatment conditions were produced for each participant using z-transformed fALFF (zfALFF) maps and statistical parametric mapping software (SPM12, Wellcome Department of Imaging Neuroscience, London, UK; http://www.fil.ion.ucl.ac.uk/spm/software/spm12). Subsequently, we employed a voxel-wise one-sample t test to examine general effects of treatment and a two-sample t test examine interactive effects between treatment (coded in the individual contrast map as within-subject factor) and GG and TT genotype (coded as between-subject factor group) within SPM12. Results that passed p < 0.05 at a cluster-level family wise error (FWE) threshold in combination with an initial threshold of p < 0.001 uncorrected at the voxel level within our search mask were reported.

### Functional connectivity analysis

Based on the results of the fALFF analysis, we calculated a seed-based voxel-wise functional connectivity matrix performed using the User-Friendly Functional Connectivity software (UF_2_C, https://www.lniunicamp.com/uf2c), which was developed by (72) and implemented in MATLAB. Specifically, a 6mm-radius sphere centered on the MNI peak coordinates of the identified effects on spontaneous neural activity served as a seed region and we examined positive and negative FC by extracting an average time-series from the seed and subsequently calculating the Pearson’s correlation coefficient between the time-series of the seed and each voxel in whole brain to generate correlation maps. The contrast maps and statistical analyses of z-transformed positive and negative FC compared between ATD and PLC treatment were separately performed as described above in fALFF analyses section. Similarly, results were thresholed at a cluster-level of p < 0.05 with family-wise error (FWE) after small volume correction (an initial voxel-level threshold at p< 0.001).

### Connectome gradient analysis

To further explore the effects of ATD and genotype on the hierarchy of cortical organization on the whole-brain level, we conducted a connectome gradient analysis (73). We first resampled the preprocessed brain images to 4×4×4 mm isotropic resolution, and then calculated the voxel-wise connectivity matrix between 18933 nodes and constructed the connectome gradient by applying diffusion map embedding approach for each subject to capture the top 10 gradients. Next, three global metrics, including gradient explanation ratio, gradient range and gradient variation, were generated for all subjects. A general linear model with age as a covariate was employed to explore the treatment and genotype group differences. The results for the global metrics were reported at a statistical significance threshold of p < 0.05. In line with previous gradient studies (73), the results for regional gradient score maps were thresholded at a voxel level of p < 0.001 with Gaussian random field (GRF) correction at the cluster level of p < 0.05.

## Results

### Sample characteristics and potential confounders

The genotype samples exhibited comparable age, stress (CTQ, PSS) and pre-treatment anxiety levels (STAI) suggesting that the TT and GG carriers in the present study were matched in terms of demographics and pre-treatment emotional state (see **Table 1**).

**Table 1.** Sample demographics and potential confounders.

|  | TT (n=23) | GG(n=25) | T, p |
| --- | --- | --- | --- |
| Age | 21.87±2.49 | 22.00±2.72 | -0.17,0.86 |
| CTQ | 40.00±8.67 | 37.76±4.84 | 1.09,0.28 |
| PSS | 13.35±5.31 | 14.48±5.61 | -0.72,0.48 |
| TAI | 38.43±8.41 | 36.72±8.53 | 0.70,0.49 |
| SAI<sub>ATD</sub> | 34.00±8.43 | 34.72±9.18 | -0.286,0.78 |
| SAI<sub>PLC</sub> | 34.83±7.82 | 32.52±7.38 | 1.05,0.30 |

Abbreviation: CTQ, Childhood Trauma Questionnaire; PSS, Perceived Stress Scale; TAI, Trait Anxiety Inventory; SAI, State Anxiety Inventory; SAI<sub>ATD</sub>, pre-treatment assessment during acute tryptophan depletion session; SAI<sub>PLC</sub>, pre-treatment assessment during placebo session.

### The effects of the reduction of 5-HT signaling via ATD on spontaneous neural activity

FALFF analysis was performed to precisely map the effects of ATD on regional intrinsic neural activity. A comparison between ATD and PLC treatment revealed a significant main effect of treatment on the intrinsic activity of the left ventral hippocampus (T= 4.47, k = 6, p_FWE_ < 0.05 within the vmPFC-amygdala-hippocampus ROI, peak MNI, x, y, z = -33, -21, -18, **Fig. 1**), reflecting that compared to placebo transient reduction in 5-HT signaling via ATD enhanced spontaneous neural activity in this region. To facilitate a more specific localization of the effects, we utilized probabilistic maps from the Anatomy2.2b toolbox which revealed that the effect was specifically located in the CA1 region of hippocampus (**Fig. 1**). No interactive effects between treatment and genotype were found.

**Fig 1.**
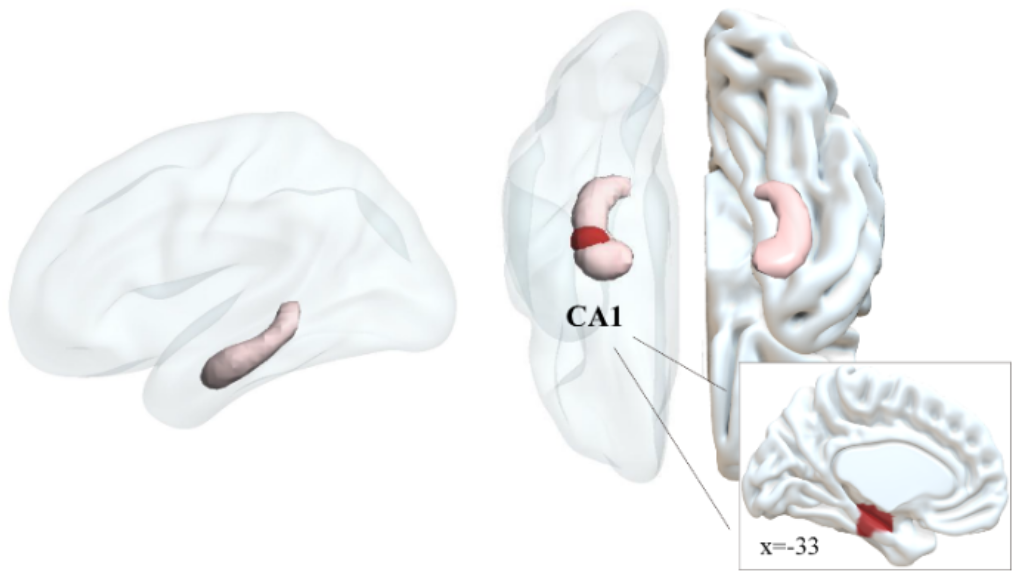
Effects of tryptophan depletion on voxel-wise spontaneous brain activity. Left plot displays the anatomical location of the hippocampus, and the right plot presents the localization of the effects of ATD-induced increased 5-HT signaling on spontaneous activity in CA1 region of left hippocampus (for display purpose shown at p < 0.005).

### Functional connectivity changes for main effects of ATD and its interaction with genotype

Based on the results of treatment effects on intrinsic activity, we quantified the functional connectivity matrix between the identified CA1 region (6-mm radius sphere centered at the peak coordinates x, y, z = -33, -21, -18) and the whole brain. We initially examined the functional connectivity of this region by pooling the data from the treatment sessions, reflecting that the region exhibited widespread connectivity with the bilateral fronto-parietal network, limbic network and sensorimotor network (displayed as **Fig. 2a**). Examining the main effects of treatment on the local network level revealed that ATD significantly decreased positive FC between the CA1 medial frontal regions, including medial frontal and orbitofrontal regions as well as further cingulate regions (**Fig. 2b**). A significant interaction effect between genotype and ATD was determined on the positive connectivity of the CA1 with the bilateral amygdala spreading into the parahippocampal gyrus, as well as vmPFC regions (**Fig. 2c**). We next extracted parameter estimates from significant regions to explore the complex interaction effect in SPSS22.0 (Fig. 2d). Bonferroni corrected post-hoc analyses on the extracted parameter estimates from significant regions revealed that in GG carriers, ATD treatment significantly enhanced functional connectivity between the CA1 with the amygdala and vmPFC (p < 0.001), while in the TT carriers ATD treatment decreased the connectivity between these regions (p = 0.047).

**Fig 2.**
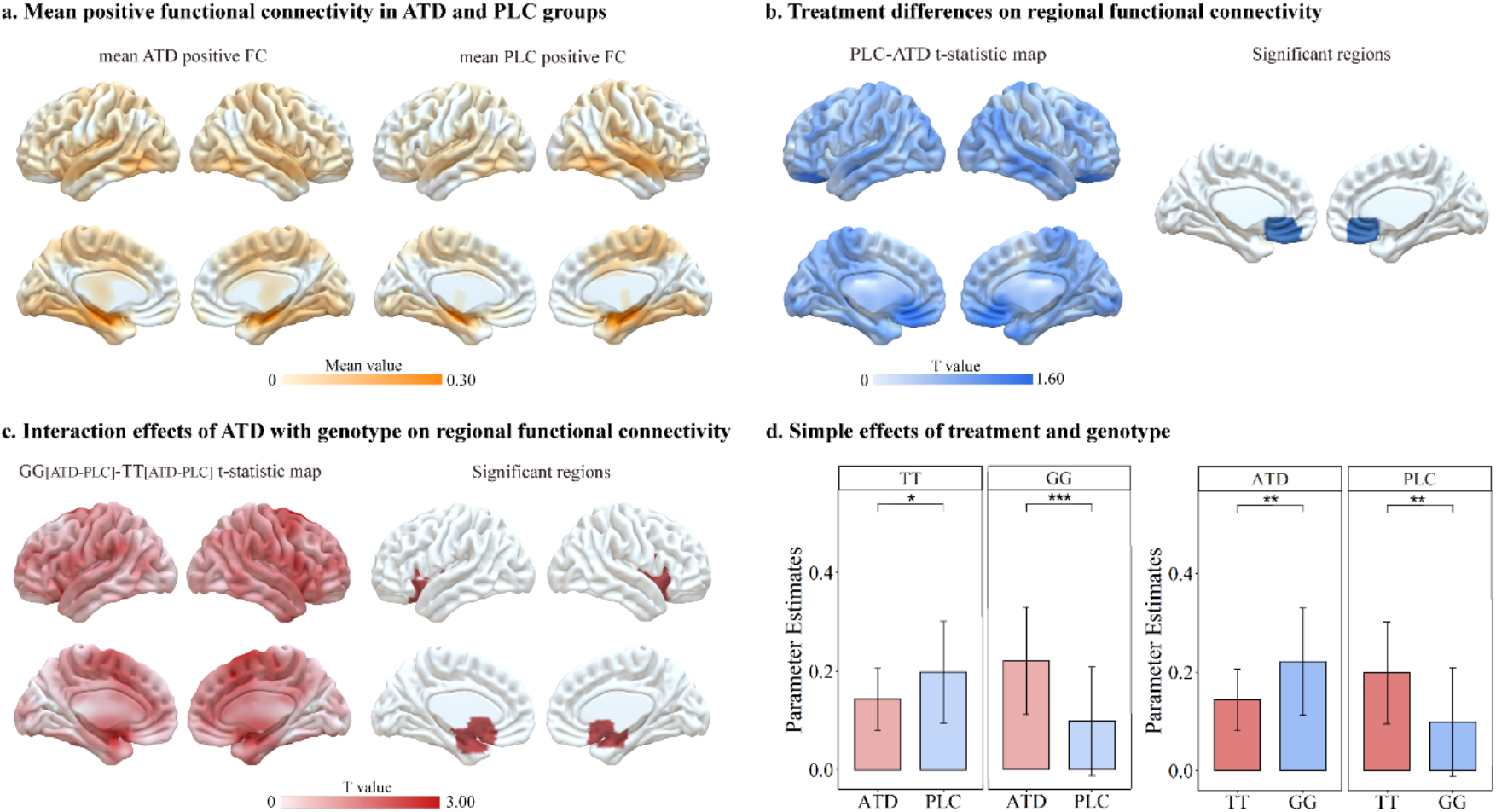
Effects of ATD treatment and its interaction effects with genotype on local functional connectivity. (for display purpose shown at p < 0.005). **(a)** Regional mean positive FC maps in the ATD and placebo groups, respectively. **(b)** Statistical t-map between ATD and PLC group, and regional alterations in positive FC induced by treatment (T = 4.10, k = 59, p_FWE_ = 0.021, peak MNI, x, y, z = 0, 30, -15). **(c)** Changes of positive functional coupling for interaction effects of treatment with genotype (cluster 1, T = 4.19, k = 23, p_FWE_ = 0.017, peak MNI, x, y, z = 18, 0, -15; cluster 2, T = 4.09, k = 48, p_FWE_ = 0.023, peak MNI, x, y, z = -24, 3, -18). **(d)** Examination of simple effects for interaction effects with extracted parameter estimates form significant regions. * indicates p < 0.05, ** indicates p < 0.01, *** indicates p < 0.001.

### ATD-induced alterations in connectome gradients on the large-scale network level

We capitalized on the diffusion map embedding method to capture connectome gradient patterns in both groups. The principal gradient 3 was represented along a gradual axis from sensorimotor network (SMN) to frontoparietal network (FPN) in both groups (**Fig. 3a, 3c**). There were some treatment differences on global topology of gradient 3, manifested on the gradient range (p = 0.04, Cohen’s d = 0.44) and gradient variance (p = 0.02, Cohen’s d = 0.54, **Fig. 3b**), however, these differences did not pass

**Fig 3.**
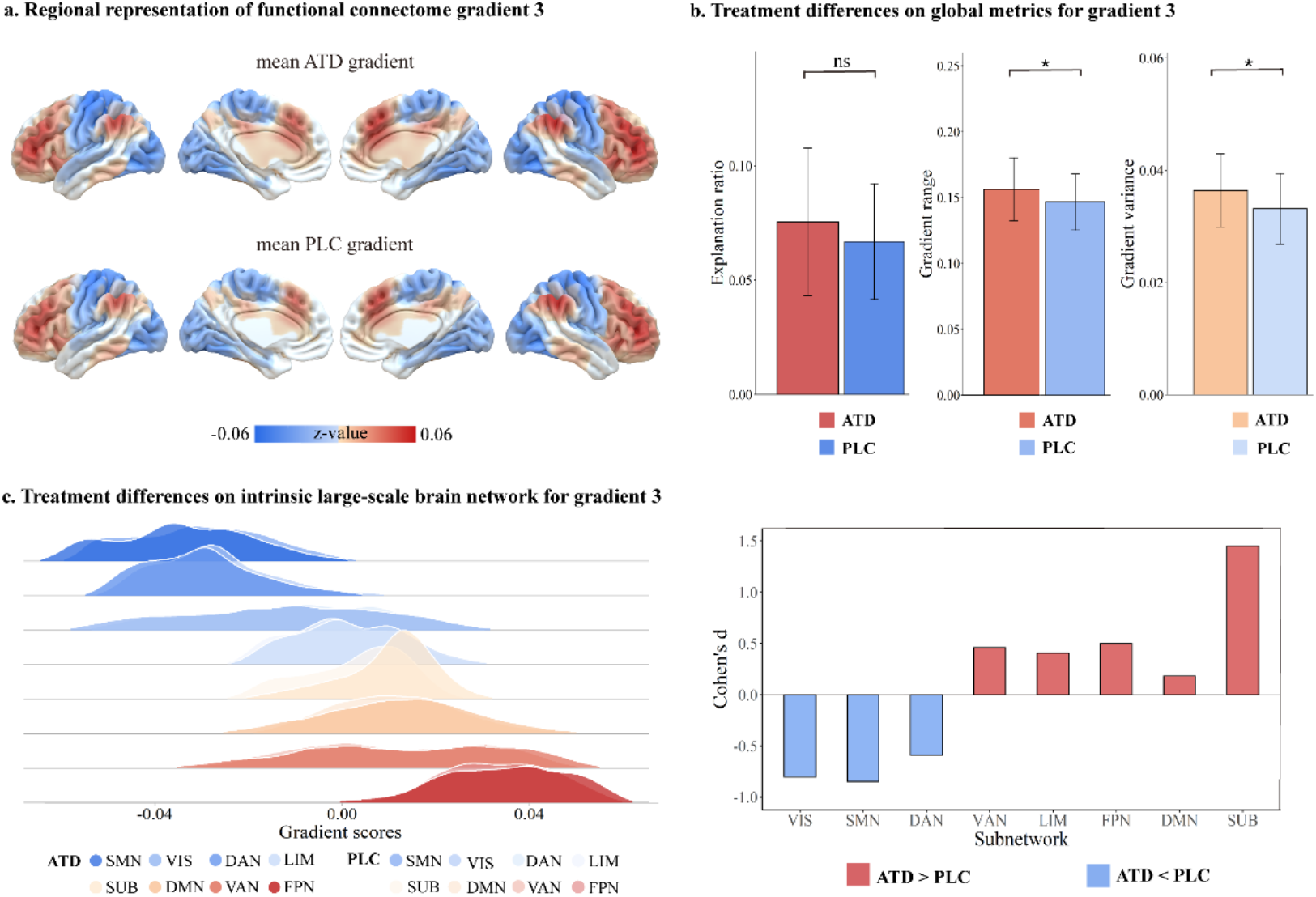
Connectome gradient mapping in ATD and placebo groups. **(a)** averaged-group gradient maps for the third gradient in ATD and PLC. **(b)** Statistics for global metrics, i.e. explanation ratio, gradient range, gradient variance of connectome gradient3. **(c)** treatment-induced differences in intrinsic large-scale brain network for gradient3. The left ridgeline plot showed the differences of density distribution for gradient scores from eight subnetworks in ATD and PLC groups, all of which met the statistical significance level and survived the Bonferroni FDR correction (all ps< 0.0001). The right bar plot depicts the effect size Cohen’s d for the difference between ATD and PLC groups in eight subnetwork systems. * indicates p< 0.05, ns indicates no significant. SMN, sensorimotor network; VIS, visual network; DAN, dorsal attention network; LIM, limbic network; SUB, subcortical regions; DMN, default mode network; VAN, ventral attention network; FPN, frontoparietal network.

Bonferroni FDR correction. In gradient 3, we found moreover a robust effect of ATD treatment on the intrinsic large-scale networks, specifically, following ATD (compared to PLC) treatment significant higher gradient scores were observed in the ventral attention network (VAN or salience network, p < 0.0001, Cohen’s d = 0.46), limbic network (LIM, p < 0.0001, Cohen’s d = 0.41), FPN (p < 0.0001, Cohen’s d = 0.50), default mode network (DMN, p < 0.0001, Cohen’s d = 0.18), subcortical network (SUB, p < 0.0001, Cohen’s d = 1.45), while lower gradient scores were observed in the visual network (VIS, p < 0.0001, Cohen’s d = -0.80), SMN (p < 0.0001, Cohen’s d = 0.85), dorsal attention network (DAN, p < 0.0001 Cohen’s d = -0.59) in ATD group (**Fig. 3c**). No other significant main effects of genotype and its interaction with treatment were found.

## Discussion

The present study aimed at determining whether individual variations in genetic polymorphisms directly related to individual variations in central serotonin synthesis rates (TPH2) influence the susceptibility to the effects of a transient decrease in 5-HT signaling on the intrinsic neural activity within limbic-prefrontal circuits, their connectivity and the whole brain gradient architecture of the brain. To this end, we recruited extreme genotype groups of a prominent TPH2 variation that underwent a pre-registered within-subject placebo-controlled pharmacological fMRI trial. Results revealed significant effects of ATD on spontaneous neural activity in hippocampal CA1 and functional coupling with vmPFC, hippocampus and amygdala. When accounting for the genetic differences, significant interaction effects between TPH2 genotype and ATD were observed, such that ATD specifically increased FC between hippocampus, amygdala and vmPFC in GG carriers, while it decreased FC in this pathway in the TT carriers. Finally, the current study, for the first time, revealed the effects of ATD on the brain internal gradient architecture, particularly in the third gradient, highlighting the role of 5-HT system in modulating the large-scale networks.

The effects of the reduction of 5-HT signaling via ATD on spontaneous neural activity specifically affected the hippocampal CA1 region which exhibits dense serotonergic innervation (74). Previous studies have reported effects of transient decreases in 5-HT signaling on the hippocampus during memory formation (75) and the processing of emotional stimuli (76) with effects being modulated by trait threat sensitivity of the participants. Studies on the effects of long-term modulation of 5-HT signaling by the use of selective serotonin inhibitors (SSRIs) reported effects in hippocampal neurogenesis in response to changes in serotonin system (77, 78, 79). Previous animal studies have elucidated that the genetic ablation of the 5-HT1A receptor precipitates alterations in synaptic activity and signal transduction, particularly within the hippocampal CA1 region promoting augmented neuronal excitability (80, 81). The 5-HT1A receptor serves as an inhibitory autoreceptor, rendering it susceptible to stress and responsive to antidepressants (82), and is implicated in the modulation of hedonic processes (83). ATD, which precipitates a reduction in serotonergic signaling, attenuates the activation of 5-HT1A receptors (84), and may thereby potentiate the spontaneous activity of hippocampal CA1 neurons. Overall, the present findings suggest robust and regional-specific effects of 5-HT modulation on spontaneous regional activity in the hippocampus, while no effects in the amygdala and vmPFC or modulatory influences of genotype were observed.

Examining effects on the network-level intrinsic communication of the identified hippocampal region revealed that ATD decreased the positive functional connectivity between this region with bilateral medial frontal regions encompassing medial frontal, orbitofrontal and anterior cingulate regions, underscoring the modulatory role of 5-HT on the communication between hippocampal and frontal regions (85) including contexts that require emotion regulation (22, 86). Interaction effects depending on the TPH2 gene polymorphism were observed on the functional connectivity of the hippocampus region with the bilateral amygdala extending into the parahippocampal gyrus as well as posterior vmPFC regions, an effect that was primarily driven by marked increased in functional connectivity in the GG carriers. These findings may reflect a heighted sensitivity to the decreased central serotonin induced by ATD in GG carriers, which partly have contributed to the lack of robust effects of ATD on neural processing in recent large-scale meta-analyses (34, 35). The hippocampus, amygdala, and prefrontal cortex are intrinsically strongly connected and together mediate critical domains (87, 88). The amygdala and hippocampus are commonly recruited during encountering potential threats, eliciting preliminary anxiety and stress responses (89, 90), with CA1-amygdala projections contributing to contextual fear encoding and renewal (91), and the further evaluation of perceived threat with contextual and mnemonic contents (92). The amygdala and hippocampus together communicate with the medial prefrontal cortex, which establishes value (93) and emotion regulation (94) and inhibits excessive amygdala and hippocampus in anxiety-related behaviors (75, 95). This flow of information allows the prefrontal cortex to modulate emotion regulation by influencing the hippocampus and amygdala, which then convey affective information back to the prefrontal cortex, thus maintaining a dynamic interplay between these brain regions, which establishes a feedback loop that underlies bottom-up responses to stress and anxiety (96). Studies support the view that enhanced attentional resources allocated to negative stimuli could be the consequence of bottom-up processes via increased connectivity in amygdala, hippocampus and prefrontal cortex, resulting in a heightened anxiety response (89, 97), similar to the observations in individuals with mood disorders, such as anxiety disorder and posttraumatic stress disorder (PTSD) (98, 99, 100, 101). The observed increased connectivity between the hippocampus, amygdala, and vmPFC after transient dysfunction in serotoninergic system thus may signal modulation in a bottom-up anxiety signaling system.

Further, this bottom-up 5-HT sensitive anxiety-related pathway was susceptible to genetic variations in the TPH2 polymorphisms which has been commonly linked to mood and anxiety disorders (102). In line with our recent study (4), GG carriers were more sensitive to the effects of transient decrease in central 5-HT signaling. The G-allele of rs4570625 SNP has been proposed to correlate with reduced activity of tryptophan hydroxylase (103), potentially leading to decreased synthesis rates of serotonin (5-HT), which could amplify the effects of temporary decreases in tryptophan levels. A growing number of studies demonstrates that the GG genotype could be associated with impaired inhibitory processes (39), which is an essential component of emotion regulation systems and critically relies on the integrity of the medial prefrontal-limbic circuits (104). The observed genotype-specific effects on hippocampus-vmPFC-amygdala coupling induced by tryptophan availability in the GG carriers may indicate that genetic predispositions can significantly influence how serotonin depletion impacts neural connectivity.

On the large-scale network level, we further revealed that transient decrease in central 5-HT signaling induced alterations in the brain’s internal gradient architecture. ATD administration enhanced gradient score in the VAN (also referred to as salience network), FPN, and DMN. Core regions of these networks have been associated with emotional reactivity and emotion regulation. Effects on the gradient architecture of the frontoparietal cortex were pronounced, with this network being critically involved in range of executive domains, including inhibitory control and emotion regulation, as well as the experience of affective arousal (105, 106, 107, 108, 109). Variations in the intrinsic architecture of the FPN have been associated with the allocation of attentional resources towards threat-like information, potentially heightening susceptibility to exacerbated negative emotion (4, 110, 111), while individuals with anxiety-related disorders exhibit a bias towards negative stimulus in attentional processes (112, 113). The effects of transient tryptophan depletion additionally encompassed the limbic network with regions such as the amygdala and the vmPFC, as well as the subcortical system including the hippocampus (114). The observations underscore the pivotal role of 5-HT, as a critical neural regulator which is strongly involved in network modulation and communication across the entire brain. However, we only observed the effect of ATD in the third gradient, possibly due to the relatively small sample size of participants, as compared to previous studies utilizing larger datasets (115, 116). On the other hand, this may underlie the diverse serotonergic functions from response to early pressure (12) and stress (13) to cognitive regulation, external attention and motor control (8, 117). Thus, our results provide a potential linkage between connectome gradient disruption and impairments in 5-HT system.

Findings of the present study need to be considered in the context of the following limitations. First, the current study was conducted exclusively with male participants, to reduce variance in the data related to sex differences in the 5-HT synthesis (118), which limits the generalizability of the findings to the broader population, including females. Future research should include a more diverse sample to determine whether the observed effects extend across genders. Second, although the ROI analysis facilitated a more robust determination of specific effects, this method focusing on pre-defined regions may not capture the full extent of brain activity changes induced by ATD and genetic variations (119). Future studies should incorporate whole-brain analyses to provide a more comprehensive view of the effects of validation and replication. Third, while our study concentrated on the TPH2 polymorphism, it did not reveal a significant impact of TPH2 on the brain internal gradient architecture, which may be due to other genetic factors or molecular mechanisms influencing 5-HT signaling, such as GABAergic transmission (120). Future research should explore additional genes involved in serotonin synthesis, transport, and receptor function. Incorporating molecular and biochemical assays to directly measure 5-HT levels and receptor activity could provide deeper insights into how genetic variations impact serotonergic function.

In conclusion, the present study provides compelling evidence of the pivotal role of 5-HT in modulating intrinsic neural activity and connectivity within specific circuits closely related to anxiety regulation, with the individual variations in a TPH2 polymorphism altering susceptibility to the neural effects of transient serotonin depletion induced by ATD. This may account for the previously inconsistent effects of ATD, thereby identifying a potential genetic marker of vulnerability for anxiety-related conditions. Moreover, our study provides the first evidence of 5-HT’s impact on brain gradient architecture, underscoring its essential role in modulating network communication across the entire brain and offering a plausible explanation for the extensive functional repertoire of serotonin. Future research should incorporate more diverse samples and whole-brain analyses to corroborate and extend these findings, as well as explore additional genetic factors that may modulate serotonergic function.

## Acknowledgments

The work was partly supported by the China MOST2030Brain Project (grant no. 2022ZD0208500), National Natural Science Foundation of China (Grants No. 32250610208, 82271583) and a start-up funding from The University of Hong Kong.

The funders had no further role in study design; in the collection, analysis and interpretation of data; in the writing of the report; and in the decision to submit the paper for publication.

LW, CL, TX and BB concepted and designed the study. LW, CL, TX, XG, WZ conducted the experiment and collected the data. LW and TX performed data analyses and contributed to the figures. LW and BB wrote the manuscript. CM, TX, XG, KK and BB revised the manuscript draft.

## Disclosures

The authors report no biomedical financial interests or potential conflicts of interest.

## Notes

### Competing Interest Statement

The authors have declared no competing interest.

## Reference

1. Salvan P, Fonseca M, Winkler AM, Beauchamp A, Lerch JP, & Johansen-Berg H. (2023). Serotonin regulation of behavior via large-scale neuromodulation of serotonin receptor networks. Nat Neurosci, 26: 53–63.

2. Etkin A, Buchel C, & Gross JJ. (2015). The neural bases of emotion regulation. Nat Rev Neurosci, 16: 693–700.

3. Liu C, Lan C, Li K, Zhou F, Yao S, Xu L, et al. (2021). Oxytocinergic modulation of threat-specific amygdala sensitization in humans is critically mediated by serotonergic mechanisms. Biol Psychiatry Cogn Neurosci Neuroimaging, 6: 1081–1089.

4. Liu C, Li K, Fu M, Zhang Y, Sindermann C, Montag C, et al. (2023). A central serotonin regulating gene polymorphism (TPH2) determines vulnerability to acute tryptophan depletion-induced anxiety and ventromedial prefrontal threat reactivity in healthy young men. Eur Neuropsychopharmacol, 77: 24–34.

5. Luo Q, Kanen JW, Bari A, Skandali N, Langley C, Knudsen GM, et al. (2024). Comparable roles for serotonin in rats and humans for computations underlying flexible decision-making. Neuropsychopharmacology, 49: 600–608.

6. Azmitia EC, & Segal M. (1978). An autoradiographic analysis of the differential ascending projections of the dorsal and median raphe nuclei in the rat. J Comp Neurol, 179: 641–667.

7. Challis C, & Berton O. (2015). Top-down control of serotonin systems by the prefrontal cortex: a path toward restored socioemotional function in depression. ACS Chem Neurosci, 6: 1040–1054.

8. Cools R, Roberts AC, & Robbins TW. (2008). Serotoninergic regulation of emotional and behavioural control processes. Trends Cogn Sci, 12: 31–40.

9. Holmes A. (2008). Genetic variation in cortico-amygdala serotonin function and risk for stress-related disease. Neurosci Biobehav Rev, 32: 1293–1314.

10. Hornboll B, Macoveanu J, Nejad A, Rowe J, Elliott R, Knudsen GM, et al. (2018). Neuroticism predicts the impact of serotonin challenges on fear processing in subgenual anterior cingulate cortex. Sci Rep, 8: 17889.

11. Zhang X, Huettel SA, O’Dhaniel A, Guo H, & Wang L. (2019). Exploring common changes after acute mental stress and acute tryptophan depletion: Resting-state fMRI studies. J Psychiatr Res, 113: 172–180.

12. Liu C, Xu L, Li J, Zhou F, Yang X, Zheng X, et al. (2021). Serotonin and early life stress interact to shape brain architecture and anxious avoidant behavior–a TPH2 imaging genetics approach. Psychol Med, 51: 2476–2484.

13. Lan C, Liu C, Li K, Zhao Z, Yang J, Ma Y, et al. (2022). Oxytocinergic modulation of stress-associated amygdala-hippocampus pathways in humans is mediated by serotonergic mechanisms. Int J Neuropsychopharmacol, 25: 807–817.

14. Booij L, Tremblay RE, Szyf M, & Benkelfat C. (2015). Genetic and early environmental influences on the serotonin system: consequences for brain development and risk for psychopathology. J Psychiatry Neurosci, 40: 5–18.

15. van der Stelt HM, Broersen LM, Olivier B, & Westenberg HG. (2004). Effects of dietary tryptophan variations on extracellular serotonin in the dorsal hippocampus of rats. Psychopharmacology, 172: 137–144.

16. Markus CR. (2008). Dietary amino acids and brain serotonin function; implications for stress-related affective changes. Neuromolecular medicine, 10: 247–258.

17. Crockett M, Clark L, Roiser J, Robinson O, Cools R, Chase H, et al. (2012). Converging evidence for central 5-HT effects in acute tryptophan depletion. Mol Psychiatry, 17: 121–123.

18. Van Donkelaar E, Blokland A, Ferrington L, Kelly P, Steinbusch H, & Prickaerts J. (2011). Mechanism of acute tryptophan depletion: is it only serotonin? Mol Psychiatry, 16: 695–713.

19. Lindseth G, Helland B, & Caspers J. (2015). The effects of dietary tryptophan on affective disorders. Arch Psychiatr Nurs, 29: 102–107.

20. Kanen JW, Arntz FE, Yellowlees R, Cardinal RN, Price A, Christmas DM, et al. (2021). Serotonin depletion amplifies distinct human social emotions as a function of individual differences in personality. Transl Psychiatry, 11: 81.

21. Ardis T, Cahir M, Elliott J, Bell R, Reynolds G, & Cooper S. (2009). Effect of acute tryptophan depletion on noradrenaline and dopamine in the rat brain. J Psychopharmacol, 23: 51–55.

22. Passamonti L, Crockett MJ, Apergis-Schoute AM, Clark L, Rowe JB, Calder AJ, et al. (2012). Effects of acute tryptophan depletion on prefrontal-amygdala connectivity while viewing facial signals of aggression. Biol Psychiatry, 71: 36–43.

23. Van der Veen FM, Evers EA, Deutz NE, & Schmitt JA. (2007). Effects of acute tryptophan depletion on mood and facial emotion perception related brain activation and performance in healthy women with and without a family history of depression. Neuropsychopharmacology, 32: 216–224.

24. Bår K-J, Köhler S, de la Cruz F, Schumann A, Zepf FD, & Wagner G. (2020). Functional consequences of acute tryptophan depletion on raphe nuclei connectivity and network organization in healthy women. Neuroimage, 207: 116362.

25. Gothard KM. (2020). Multidimensional processing in the amygdala. Nat Rev Neurosci, 21: 565–575.

26. Mihov Y, Kendrick KM, Becker B, Zschernack J, Reich H, Maier W, et al. (2013). Mirroring fear in the absence of a functional amygdala. Biol Psychiatry, 73: e9–e11.

27. Zhou F, Geng Y, Xin F, Li J, Feng P, Liu C, et al. (2019). Human extinction learning is accelerated by an angiotensin antagonist via ventromedial prefrontal cortex and its connections with basolateral amygdala. Biol Psychiatry, 86: 910–920.

28. da Cunha-Bang S, & Knudsen GM. (2021). The modulatory role of serotonin on human impulsive aggression. Biol Psychiatry, 90: 447–457.

29. Crockett MJ, Clark L, Tabibnia G, Lieberman MD, & Robbins TW. (2008). Serotonin modulates behavioral reactions to unfairness. Science, 320: 1739–1739.

30. Crockett MJ, Clark L, Lieberman MD, Tabibnia G, & Robbins TW. (2010). Impulsive choice and altruistic punishment are correlated and increase in tandem with serotonin depletion. Emotion, 10: 855.

31. Cools R, Calder AJ, Lawrence AD, Clark L, Bullmore E, & Robbins TW. (2005). Individual differences in threat sensitivity predict serotonergic modulation of amygdala response to fearful faces. Psychopharmacology, 180: 670–679.

32. Herringa RJ, Birn RM, Ruttle PL, Burghy CA, Stodola DE, Davidson RJ, et al. (2013). Childhood maltreatment is associated with altered fear circuitry and increased internalizing symptoms by late adolescence. Proc Natl Acad Sci U S A, 110: 19119–19124.

33. Fan Y, Pestke K, Feeser M, Aust S, Pruessner JC, Böker H, et al. (2015). Amygdala–hippocampal connectivity changes during acute psychosocial stress: joint effect of early life stress and oxytocin. Neuropsychopharmacology, 40: 2736–2744.

34. Raab K, Kirsch P, & Mier D. (2016). Understanding the impact of 5-HTTLPR, antidepressants, and acute tryptophan depletion on brain activation during facial emotion processing: A review of the imaging literature. Neurosci Biobehav Rev, 71: 176–197.

35. Schopman SM, Bosman RC, Muntingh AD, van Balkom AJ, & Batelaan NM. (2021). Effects of tryptophan depletion on anxiety, a systematic review. Transl Psychiatry, 11: 118.

36. Bîlc MI, Iacob A, Szekely-Copîndean RD, Kiss B, Ştefan M-G, Mureşan RC, et al. (2023). Serotonin and emotion regulation: the impact of tryptophan depletion on emotional experience, neural and autonomic activity. Cogn Affect Behav Neurosci, 23: 1414–1427.

37. Eisner P, Klasen M, Wolf D, Zerres K, Eggermann T, Eisert A, et al. (2017). Cortico-limbic connectivity in MAOA-L carriers is vulnerable to acute tryptophan depletion. Hum Brain Mapp, 38: 1622–1635.

38. Kanen JW, Arntz FE, Yellowlees R, Christmas DM, Price A, Apergis-Schoute AM, et al. (2021). Effect of tryptophan depletion on conditioned threat memory expression: role of intolerance of uncertainty. Biological Psychiatry: Cognitive Neuroscience and Neuroimaging, 6: 590–598.

39. Booij L, Turecki G, Leyton M, Gravel P, Lopez De Lara C, Diksic M, et al. (2012). Tryptophan hydroxylase2 gene polymorphisms predict brain serotonin synthesis in the orbitofrontal cortex in humans. Mol Psychiatry, 17: 809–817.

40. Furmark T, Marteinsdottir I, Frick A, Heurling K, Tillfors M, Appel L, et al. (2016). Serotonin synthesis rate and the tryptophan hydroxylase-2: G-703T polymorphism in social anxiety disorder. J Psychopharmacol, 30: 1028–1035.

41. Gutknecht L, Jacob C, Strobel A, Kriegebaum C, Müller J, Zeng Y, et al. (2007). Tryptophan hydroxylase-2 gene variation influences personality traits and disorders related to emotional dysregulation. Int J Neuropsychopharmacol, 10: 309–320.

42. Laas K, Kiive E, Måestu J, Vaht M, Veidebaum T, & Harro J. (2017). Nice guys: Homozygocity for the TPH2-703G/T (rs4570625) minor allele promotes low aggressiveness and low anxiety. J Affect Disord, 215: 230–236.

43. Wigner P, Czarny P, Synowiec E, Bijak M, Białek K, Talarowska M, et al. (2018). Association between single nucleotide polymorphisms of TPH1 and TPH2 genes, and depressive disorders. J Cell Mol Med, 22: 1778–1791.

44. Markett S, de Reus MA, Reuter M, Montag C, Weber B, Schoene-Bake J-C, et al. (2017). Serotonin and the Brain’s Rich Club—Association Between Molecular Genetic Variation on the TPH2 Gene and the Structural Connectome. Cereb Cortex, 27: 2166–2174.

45. Deng S, Franklin CG, O’Boyle M, Zhang W, Heyl BL, Jerabek PA, et al. (2022). Hemodynamic and metabolic correspondence of resting-state voxel-based physiological metrics in healthy adults. Neuroimage, 250: 118923.

46. Xu T, Chen Z, Zhou X, Wang L, Zhou F, Yao D, et al. (2024). The central renin–angiotensin system: A genetic pathway, functional decoding, and selective target engagement characterization in humans. Proc Natl Acad Sci U S A, 121: e2306936121.

47. Nishizawa S, Benkelfat C, Young SN, Leyton M, Mzengeza S, de Montigny C, et al. (1997). Differences between males and females in rates of serotonin synthesis in human brain. Proceedings of the National Academy of Sciences of the United States of America, 94: 5308–5313.

48. Sambeth A, Blokland A, Harmer CJ, Kilkens TO, Nathan PJ, Porter RJ, et al. (2007). Sex differences in the effect of acute tryptophan depletion on declarative episodic memory: a pooled analysis of nine studies. Neurosci Biobehav Rev, 31: 516–529.

49. Ellenbogen MA, Young SN, Dean P, Palmour RM, & Benkelfat C. (1996). Mood response to acute tryptophan depletion in healthy volunteers: sex differences and temporal stability. Neuropsychopharmacology, 15: 465–474.

50. Duncan NW, Hayes DJ, Wiebking C, Tiret B, Pietruska K, Chen DQ, et al. (2015). Negative childhood experiences alter a prefrontal-insular-motor cortical network in healthy adults: A preliminary multimodal rsfMRI-fMRI-MRS-dMRI study. Hum Brain Mapp, 36: 4622–4637.

51. Hahn A, Stein P, Windischberger C, Weissenbacher A, Spindelegger C, Moser E, et al. (2011). Reduced resting-state functional connectivity between amygdala and orbitofrontal cortex in social anxiety disorder. Neuroimage, 56: 881–889.

52. Soares JM, Sampaio A, Ferreira LM, Santos NC, Marques P, Marques F, et al. (2013). Stress Impact on Resting State Brain Networks. PLoS One, 8: e66500.

53. Ansorge MS, Zhou M, Lira A, Hen R, & Gingrich JA. (2004). Early-life blockade of the 5-HT transporter alters emotional behavior in adult mice. Science, 306: 879–881.

54. Spielberger CD. (1970). Manual for the State-Trait Anxiety Inventory. Self Evaluation Questionnaire. 55.

55. Cohen S, Kamarck T, & Mermelstein R. (1893). A global measure of perceived stress. Journal of Health and Social Behavior, 24: 385–396.

56. Bernstein DP, Stein JA, Newcomb MD, Walker E, Pogge D, Ahluvalia T, et al. (2003). Development and validation of a brief screening version of the Childhood Trauma Questionnaire. Child Abuse Negl, 27: 169–190.

57. Shek DT. (1988). Reliability and factorial structure of the Chinese version of the State-Trait Anxiety Inventory. Journal of Psychopathology and Behavioral Assessment, 10: 303–317.

58. Zhao X, Zhang Y, Li L, & Zhou Y. (2005). Evaluation on reliability and validity of Chinese version of childhood trauma questionnaire. Chinese Journal of Tissue Engineering Research: 209–211.

59. Huang F, Wang H, Wang Z, Zhang J, Du W, Su C, et al. (2020). Psychometric properties of the perceived stress scale in a community sample of Chinese. BMC psychiatry, 20: 1–7.

60. Williams W, Shoaf S, Hommer D, Rawlings R, & Linnoila M. (1999). Effects of acute tryptophan depletion on plasma and cerebrospinal fluid tryptophan and 5-hydroxyindoleacetic acid in normal volunteers. J Neurochem, 72: 1641–1647.

61. Moreno FA, Parkinson D, Palmer C, Castro WL, Misiaszek J, El Khoury A, et al. (2010). CSF neurochemicals during tryptophan depletion in individuals with remitted depression and healthy controls. Eur Neuropsychopharmacol, 20: 18–24.

62. Salomon RM, Cowan RL, Rogers BP, Dietrich MS, Bauernfeind AL, Kessler RM, et al. (2011). Time series fMRI measures detect changes in pontine raphe following acute tryptophan depletion. Psychiatry Res, 191: 112–121.

63. Yatham LN, Liddle PF, Sossi V, Erez J, Vafai N, Lam RW, et al. (2012). Positron emission tomography study of the effects of tryptophan depletion on brain serotonin2 receptors in subjects recently remitted from major depression. Arch Gen Psychiatry, 69: 601–609.

64. Carpenter LL, Anderson GM, Pelton GH, Gudin JA, Kirwin PD, Price LH, et al. (1998). Tryptophan depletion during continuous CSF sampling in healthy human subjects. Neuropsychopharmacology, 19: 26–35.

65. Williams WA, Shoaf SE, Hommer D, Rawlings R, & Linnoila M. (1999). Effects of acute tryptophan depletion on plasma and cerebrospinal fluid tryptophan and 5-hydroxyindoleacetic acid in normal volunteers. J Neurochem, 72: 1641–1647.

66. Young SN, Smith SE, Pihl RO, & Ervin FR. (1985). Tryptophan depletion causes a rapid lowering of mood in normal males. Psychopharmacology 87: 173–177.

67. Wang J, Wang X, Xia M, Liao X, Evans A, & He Y. (2015). GRETNA: a graph theoretical network analysis toolbox for imaging connectomics. Front Hum Neurosci, 9: 386.

68. Xie Y, Xu Z, Xia M, Liu J, Shou X, Cui Z, et al. (2022). Alterations in Connectome Dynamics in Autism Spectrum Disorder: A Harmonized Mega- and Meta-analysis Study Using the Autism Brain Imaging Data Exchange Dataset. Biol Psychiatry, 91: 945–955.

69. Tzourio-Mazoyer N, Landeau B, Papathanassiou D, Crivello F, Etard O, Delcroix N, et al. (2002). Automated anatomical labeling of activations in SPM using a macroscopic anatomical parcellation of the MNI MRI single-subject brain. Neuroimage, 15: 273–289.

70. Zou QH, Zhu CZ, Yang Y, Zuo XN, Long XY, Cao QJ, et al. (2008). An improved approach to detection of amplitude of low-frequency fluctuation (ALFF) for resting-state fMRI: fractional ALFF. J Neurosci Methods, 172: 137–141.

71. Chen YC, Xia W, Luo B, Muthaiah VP, Xiong Z, Zhang J, et al. (2015). Frequency-specific alternations in the amplitude of low-frequency fluctuations in chronic tinnitus. Front Neural Circuits, 9: 67.

72. de Campos BM, Coan AC, Lin Yasuda C, Casseb RF, & Cendes F. (2016). Large-scale brain networks are distinctly affected in right and left mesial temporal lobe epilepsy. Hum Brain Mapp, 37: 3137–3152.

73. Xia M, Liu J, Mechelli A, Sun X, Ma Q, Wang X, et al. (2022). Connectome gradient dysfunction in major depression and its association with gene expression profiles and treatment outcomes. Mol Psychiatry, 27: 1384–1393.

74. Schneck N, Tu T, Falcone HR, Miller JM, Zanderigo F, Sublette ME, et al. (2021). Large-scale network dynamics in neural response to emotionally negative stimuli linked to serotonin 1A binding in major depressive disorder. Mol Psychiatry, 26: 2393–2401.

75. Padilla-Coreano N, Canetta S, Mikofsky RM, Alway E, Passecker J, Myroshnychenko MV, et al. (2019). Hippocampal-prefrontal theta transmission regulates avoidance behavior. Neuron, 104: 601-610. e604.

76. Fusar-Poli P, Allen P, Lee F, Surguladze S, Tunstall N, Fu CH, et al. (2007). Modulation of neural response to happy and sad faces by acute tryptophan depletion. Psychopharmacology, 193: 31–44.

77. Airan RD, Meltzer LA, Roy M, Gong Y, Chen H, & Deisseroth K. (2007). High-speed imaging reveals neurophysiological links to behavior in an animal model of depression. Science, 317: 819–823.

78. Schmuckermair C, Gaburro S, Sah A, Landgraf R, Sartori SB, & Singewald N. (2013). Behavioral and neurobiological effects of deep brain stimulation in a mouse model of high anxiety-and depression-like behavior. Neuropsychopharmacology, 38: 1234–1244.

79. Dale E, Pehrson AL, Jeyarajah T, Li Y, Leiser SC, Smagin G, et al. (2016). Effects of serotonin in the hippocampus: how SSRIs and multimodal antidepressants might regulate pyramidal cell function. CNS spectrums, 21: 143–161.

80. Gordon JA, Lacefield CO, Kentros CG, & Hen R. (2005). State-dependent alterations in hippocampal oscillations in serotonin 1A receptor-deficient mice. J Neurosci, 25: 6509–6519.

81. Freeman-Daniels E, Beck SG, & Kirby LG. (2011). Cellular correlates of anxiety in CA1 hippocampal pyramidal cells of 5-HT 1A receptor knockout mice. Psychopharmacology, 213: 453–463.

82. Richardson-Jones JW, Craige CP, Guiard BP, Stephen A, Metzger KL, Kung HF, et al. (2010). 5-HT1A autoreceptor levels determine vulnerability to stress and response to antidepressants. Neuron, 65: 40–52.

83. Hori Y, Mimura K, Nagai Y, Hori Y, Kumata K, Zhang M-R, et al. (2024). Reduced serotonergic transmission alters sensitivity to cost and reward via 5-HT1A and 5-HT1B receptors in monkeys. PLoS Biol, 22: e3002445.

84. Cahir M, Ardis T, Reynolds GP, & Cooper SJ. (2007). Acute and chronic tryptophan depletion differentially regulate central 5-HT 1A and 5-HT 2A receptor binding in the rat. Psychopharmacology, 190: 497–506.

85. Unal G, Joshi A, Viney TJ, Kis V, & Somogyi P. (2015). Synaptic targets of medial septal projections in the hippocampus and extrahippocampal cortices of the mouse. J Neurosci, 35: 15812–15826.

86. Phillips ML, Medford N, Young A, Williams L, Williams SC, Bullmore E, et al. (2001). Time courses of left and right amygdalar responses to fearful facial expressions. Hum Brain Mapp, 12: 193–202.

87. Geiller T, Sadeh S, Rolotti SV, Blockus H, Vancura B, Negrean A, et al. (2022). Local circuit amplification of spatial selectivity in the hippocampus. Nature, 601: 105–109.

88. Topolnik L, & Tamboli S. (2022). The role of inhibitory circuits in hippocampal memory processing. Nat Rev Neurosci, 23: 476–492.

89. Marcinkiewcz CA, Mazzone CM, D’Agostino G, Halladay LR, Hardaway JA, DiBerto JF, et al. (2016). Serotonin engages an anxiety and fear-promoting circuit in the extended amygdala. Nature, 537: 97–101.

90. Surget A, & Belzung C. (2022). Adult hippocampal neurogenesis shapes adaptation and improves stress response: a mechanistic and integrative perspective. Mol Psychiatry, 27: 403–421.

91. Xu C, Krabbe S, Gründemann J, Botta P, Fadok JP, Osakada F, et al. (2016). Distinct hippocampal pathways mediate dissociable roles of context in memory retrieval. Cell, 167: 961-972. e916.

92. Smith AP, Stephan KE, Rugg MD, & Dolan RJ. (2006). Task and content modulate amygdala-hippocampal connectivity in emotional retrieval. Neuron, 49: 631–638.

93. Winecoff A, Clithero JA, Carter RM, Bergman SR, Wang L, & Huettel SA. (2013). Ventromedial prefrontal cortex encodes emotional value. J Neurosci, 33: 11032–11039.

94. Motzkin JC, Philippi CL, Wolf RC, Baskaya MK, & Koenigs M. (2015). Ventromedial prefrontal cortex is critical for the regulation of amygdala activity in humans. Biol Psychiatry, 77: 276–284.

95. Parfitt GM, Nguyen R, Bang JY, Aqrabawi AJ, Tran MM, Seo DK, et al. (2017). Bidirectional control of anxiety-related behaviors in mice: role of inputs arising from the ventral hippocampus to the lateral septum and medial prefrontal cortex. Neuropsychopharmacology, 42: 1715–1728.

96. Zhang W-H, Zhang J-Y, Holmes A, & Pan B-X. (2021). Amygdala circuit substrates for stress adaptation and adversity. Biol Psychiatry, 89: 847–856.

97. Nawa NE, & Ando H. (2019). Effective connectivity within the ventromedial prefrontal cortex-hippocampus-amygdala network during the elaboration of emotional autobiographical memories. NeuroImage, 189: 316–328.

98. Morey RA, Haswell CC, Hooper SR, & De Bellis MD. (2016). Amygdala, hippocampus, and ventral medial prefrontal cortex volumes differ in maltreated youth with and without chronic posttraumatic stress disorder. Neuropsychopharmacology, 41: 791–801.

99. Marin M-F, Zsido RG, Song H, Lasko NB, Killgore WD, Rauch SL, et al. (2017). Skin conductance responses and neural activations during fear conditioning and extinction recall across anxiety disorders. JAMA psychiatry, 74: 622–631.

100. Xu X, Dai J, Chen Y, Liu C, Xin F, Zhou X, et al. (2021). Intrinsic connectivity of the prefrontal cortex and striato-limbic system respectively differentiate major depressive from generalized anxiety disorder. Neuropsychopharmacology, 46: 791–798.

101. Liu Q, Zhou B, Zhang X, Qing P, Zhou X, Zhou F, et al. (2023). Abnormal multi-layered dynamic cortico-subcortical functional connectivity in major depressive disorder and generalized anxiety disorder. Journal of Psychiatric Research, 167: 23–31.

102. Gao J, Pan Z, Jiao Z, Li F, Zhao G, Wei Q, et al. (2012). TPH2 gene polymorphisms and major depression–a meta-analysis. PloS One, 7: e36721.

103. Lin Y-MJ, Chao S-C, Chen T-M, Lai T-J, Chen J-S, & Sun HS. (2007). Association of functional polymorphisms of the human tryptophan hydroxylase 2 gene with risk for bipolar disorder in Han Chinese. Archives of general psychiatry, 64: 1015–1024.

104. Gross JJ. (2013). Emotion regulation: taking stock and moving forward. Emotion, 13: 359.

105. Capotosto P, Babiloni C, Romani GL, & Corbetta M. (2009). Frontoparietal cortex controls spatial attention through modulation of anticipatory alpha rhythms. Journal of Neuroscience, 29: 5863–5872.

106. Scolari M, Seidl-Rathkopf KN, & Kastner S. (2015). Functions of the human frontoparietal attention network: Evidence from neuroimaging. Current opinion in behavioral sciences, 1: 32–39.

107. Zhuang Q, Qiao L, Xu L, Yao S, Chen S, Zheng X, et al. (2023). The right inferior frontal gyrus as pivotal node and effective regulator of the basal ganglia-thalamocortical response inhibition circuit. Psychoradiology, 3: kkad016.

108. Zhuang Q, Xu L, Zhou F, Yao S, Zheng X, Zhou X, et al. (2021). Segregating domain-general from emotional context-specific inhibitory control systems-ventral striatum and orbitofrontal cortex serve as emotion-cognition integration hubs. Neuroimage, 238: 118269.

109. Liu X, Jiao G, Zhou F, Kendrick KM, Yao D, Gong Q, et al. (2024). A neural signature for the subjective experience of threat anticipation under uncertainty. Nature Communications, 15: 1544.

110. Dainer-Best J, Trujillo LT, Schnyer DM, & Beevers CG. (2017). Sustained engagement of attention is associated with increased negative self-referent processing in major depressive disorder. Biological Psychology, 129: 231–241.

111. Xin F, Zhou F, Zhou X, Ma X, Geng Y, Zhao W, et al. (2021). Oxytocin modulates the intrinsic dynamics between attention-related large-scale networks. Cerebral Cortex, 31: 1848–1860.

112. Bar-Haim Y, Lamy D, Pergamin L, Bakermans-Kranenburg MJ, & Van Ijzendoorn MH. (2007). Threat-related attentional bias in anxious and nonanxious individuals: a meta-analytic study. Psychological bulletin, 133: 1.

113. Nejati V, Khalaji S, Goodarzi H, & Nitsche M. (2021). The role of ventromedial and dorsolateral prefrontal cortex in attention and interpretation biases in individuals with general anxiety disorder (GAD): a tDCS study. Journal of Psychiatric Research, 144: 269–277.

114. Xia M, Liu J, Mechelli A, Sun X, Ma Q, Wang X, et al. (2022). Connectome gradient dysfunction in major depression and its association with gene expression profiles and treatment outcomes. Mol Psychiatry, 27: 1384–1393.

115. Xia Y, Xia M, Liu J, Liao X, Lei T, Liang X, et al. (2022). Development of functional connectome gradients during childhood and adolescence. Sci Bull, 67: 1049–1061.

116. Sun X, Sun J, Lu X, Dong Q, Zhang L, Wang W, et al. (2023). Mapping neurophysiological subtypes of major depressive disorder using normative models of the functional connectome. Biol Psychiatry, 94: 936–947.

117. Quist J, Barr C, Schachar R, Roberts W, Malone M, Tannock R, et al. (2003). The serotonin 5-HT1B receptor gene and attention deficit hyperactivity disorder. Mol Psychiatry, 8: 98–102.

118. Nishizawa S, Benkelfat C, Young S, Leyton M, Mzengeza Sd, De Montigny C, et al. (1997). Differences between males and females in rates of serotonin synthesis in human brain. Proceedings of the national academy of sciences, 94: 5308–5313.

119. Cremers HR, Wager TD, & Yarkoni T. (2017). The relation between statistical power and inference in fMRI. PloS one, 12: e0184923.

120. Andolina D, Maran D, Valzania A, Conversi D, & Puglisi-Allegra S. (2013). Prefrontal/amygdalar system determines stress coping behavior through 5-HT/GABA connection. Neuropsychopharmacology, 38: 2057–2067.

